# A Multiscale Model for Quantitative Prediction of Insulin Aggregation Nucleation Kinetics

**DOI:** 10.1101/2021.02.13.431119

**Authors:** Rit Pratik Mishra, Gaurav Goel

## Abstract

We combined kinetic, thermodynamic, and structural information from single molecule (protein folding) and two molecule (association) explicit-solvent simulations for determination of kinetic parameters in protein aggregation nucleation with insulin as model protein. A structural bioinformatics approach was developed to account for heterogeneity of aggregation-prone species with the transition complex theory found applicable in modeling association kinetics involving non-native species. We show that a key simplification arises from presence of only a few relevant modes for non-native association kinetics. The kinetic parameters thus obtained were used in a population balance model and accurate predictions for aggregation nucleation time varying over two orders of magnitude with changes in concentration of insulin or an aggregation-inhibitor ligand were obtained while an empirical parameter set was not found to be transferable for prediction of ligand effects. This physically determined kinetic parameter set also allowed identification of the rate-limiting step in aggregation nucleation.

**TOC Graphic:** 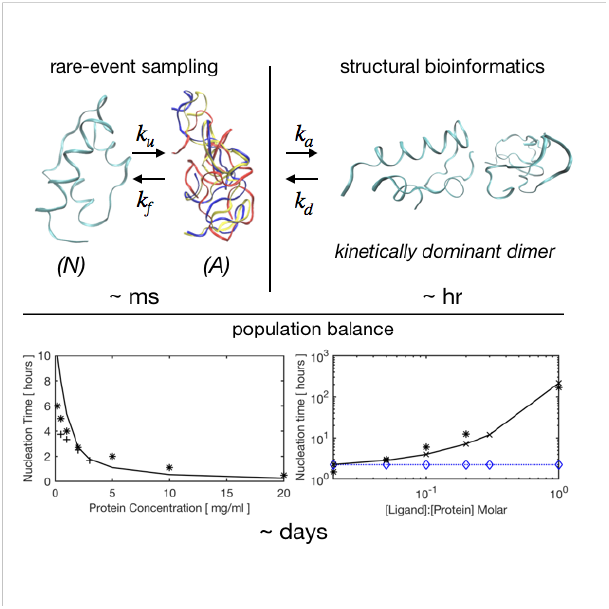

---

Protein drugs are a very important class of biotherapeutics with over ten thousands marketed products. Their in vivo or in vitro aggregation leads to issues such as loss of efficacy during storage^1^ or several severe disorders and diseases.^2^ A molecular level understanding of aggregation pathways can help in devising mitigation strategies. In particular, the knowledge of thermodynamics and kinetics of early stage aggregation nucleation events is necessary for developing strategies for inhibition of small oligomer formation that have been shown to be more cytotoxic than mature fibrils.^3^ However, their structural characterization^4,5^ and measurement of kinetic rate constants^6–8^ by bioanalytical techniques is made difficult by short life-time and a distinct, protein-dependent structure of species involved.The disparate timescales and lengthscales involved even for small proteins (< 12 kDa)—milliseconds and nanometers for protein folding, seconds and tens of nanometers for aggregation nucleation, and hours–days and centimeters for fibrillation, preclude use of all-atom explicit solvent simulations for direct determination of aggregation thermodynamics and kinetics. Solvent-free, residue specific interaction models make accessible these timescales and lengthscales.^9,10^ However, these are biased towards formation of native interactions, and hence, not suitable for studying protein association leading to aggregation.

In another set of coarsegrained techniques, a protein is modeled as an isotropic or a patchy colloidal particle and protein association kinetics are then studied using either mesoscale simulations (Brownian dynamics, dissipative particle dynamics) or event-based methods (kinetic Monte Carlo simulations, population balance modeling (PBM)). In several applications of these event-based approaches, the rates associated with elementary aggregation pathway steps are determined empirically to reproduce experimental measures of aggregation such as nuclei size, nucleation time, and growth kinetics.^11^ For example, PBM has been used to study the effect of osmolytes, protein concentration, stress factors like temperature, pH, salt, impurities, and mechanical perturbation on protein aggregation.^12–16^ In this work, we present an approach for determination of a set of physically relevant kinetic parameters and show that a single parameter set is sufficient for accurate extrapolation of aggregation behaviour on variation of multiple factors.Insulin was used as a model protein because of its biotherapeutic relevance and availability of a large number of experimental studies on advancing understanding of its aggregation mechanism and kinetics.^17,18^

The extended Lumry-Eyring model,^19^ used here to model aggregation kinetics, has been shown to be applicable to a large number of globular proteins^11^ including insulin.^12,16^ The overall model scheme, shown in Table 1, was divided in two parts: formation of a critical nucleus consisting of 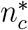 monomers by reversible addition of an aggregation-prone monomeric species (*A*) to other monomers and small oligomers (**Scheme A**, equations 1–3) followed by an aggregation growth stage involving irreversible association and, possibly, conformational rearrangement of stable oligomers (size 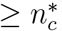) to form fibrils (**Scheme D**, equations 8– 9). The first stage, referred to as early events of aggregation, is of interest in this work. It has been shown to have significant effect on aggregation rates and, in fact, is the rate-determining step for many small proteins including insulin.^19^ In this case, the nucleation kinetics are determined by two key events: single molecule reconfiguration characterized by protein folding (*k*_f_) and unfolding (*k*_u_) rates, and dimer and higher oligomer formation characterized by protein-protein association (*k*_nu_) and dissociation (*k*_−nu_) rates. These kinetic constants together determine the nucleation time (*t*_nu_), defined as the induction time at which a measurable amount of stable oligomeric species have formed. An alternate scheme for aggregation nucleation that explicitly represents formation of a diffusional intermediate (involving a negligible change in conformation of associating species) followed by conformational rearrangement is shown in **Scheme B** (equations 4–6). A competing pathway involving aggregation inhibition by a ligand (*L*) having a high affinity for aggregation-prone species (*A*_*i*_) is shown in **Scheme C** (equation 7). Role of both of these schemes in modulating aggregation kinetics and determination of kinetic constants appearing within are discussed later in this letter. We note that it is extremely important to account for the specificity in protein-protein and protein-ligand interactions involving non-native aggregation-prone species and their effect on protein dimerization kinetics, the latter being a critical input to our model. To this end, we based our development on extensive explicit solvent simulations of a system consisting of either a single solute molecule (protein or ligand) or a single complex (protein dimer or protein-ligand complex) in-combination with a conformational selection model for protein association.^20,21^ Protein binding and folding in crowded environment, e.g. in cells, has been shown to occur at rates close to those determined in dilute solutions,^22^ implying that data obtained from simulations in the infinite dilution limit, i.e., only one (for protein folding) or two protein molecules (for protein dimerization) solvated in a large simulation box, will be relevant for modeling of aggregation kinetics.

**Table 1:**
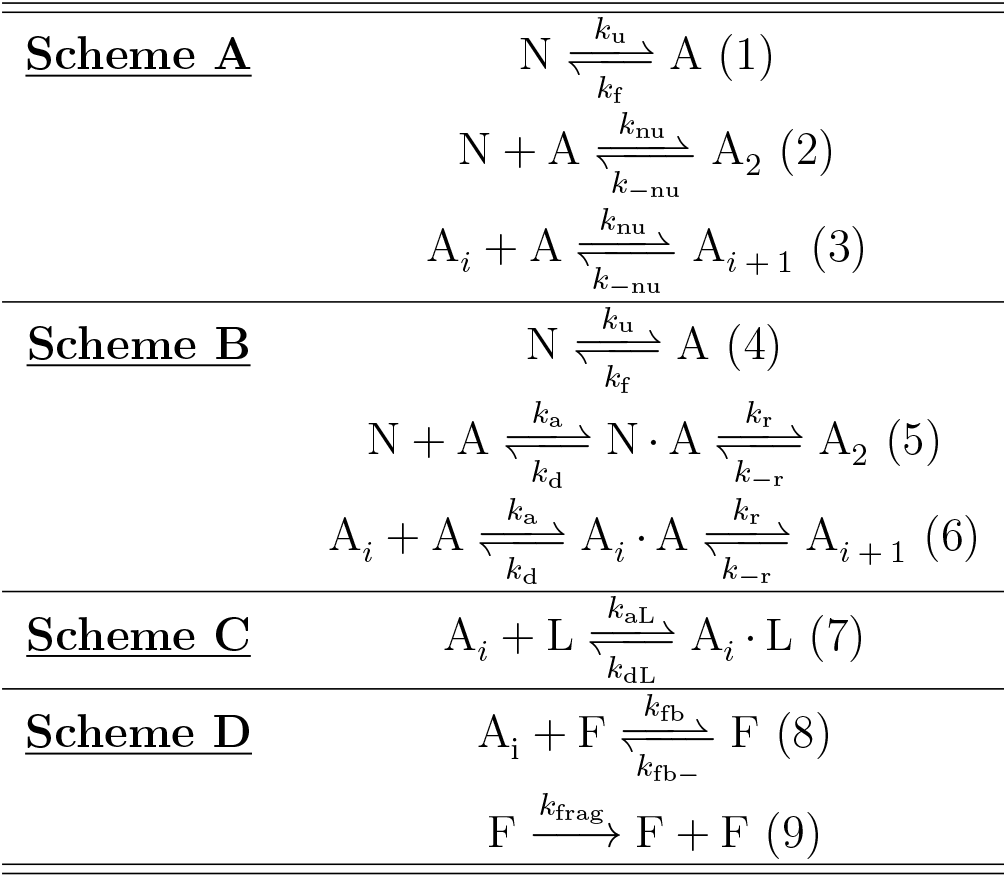
Aggregation kinetics schemes. Scheme A: early events of aggregation that include protein folding-unfolding (rates: *k*_u_, *k*_f_) and sequential addition of an aggregation-prone monomeric species (*A*_1_, also referred to as *A* for ease of notation) (rates: *k*_nu_, *k*_−nu_) to form higher order oligomeric species (*A*_*i*_) up to a stable nuclei of size 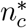 (critical nucleus). Scheme B: same as Scheme A, but the protein association step explicitly represents diffusional association (rates: *k*_a_, *k*_d_) and conformational rearrangement (rates: *k*_r_, *k*_−r_). Scheme C: addition of ligand to aggregation-prone species smaller than the critical nucleus (rates: *k*_aL_, *k*_dL_). Scheme D: formation of higher-order fibrillar species and fibril fragmentation (rates: *k*_fib_, *k*_fib−_, *k*_frag_. For all schemes, 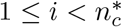. All species of size 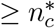 are labelled *F*.

## Reduced Kinetic Model

Explicit solvent simulations to model protein association in large oligomer and fibrillar assemblies are intractable. We, therefore, investigated the effect of excluding fibrillation steps on species concentration and on estimation of the aggregation nucleation time (*t*_nu_) of insulin. A critical nucleus size of 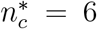 was taken in accordance with earlier experimental and computational studies on insulin aggregation.^12,16,23,24^ We calculated monomer (*C*_1_) and dimer (*C*_2_) concentrations (Figure 1) and the stable oligomer (*C*_*i*≥6_ ≡*C*_F_) concentration (Figure 3A) by solving either the full kinetic model (**Scheme A** and **Scheme D**) or only the early events kinetic model (**Scheme A**). The kinetic parameters have been taken from an earlier study wherein the full kinetic model was fitted to data on insulin aggregation growth.^12^ The association (*k*_nu_) and dissociation (*k*_−nu_) rates for addition of species *A* to any non-fibrillar species have been taken to be independent of oligomer size. The former can be explained on the basis of cancellation of the opposing effect on number of aggregation hotspots (increase with oligomer size) and oligomer rotational and translational diffusivity (decrease with oligomer size). The association rate for addition of *A* to a pentameric oligomer is only two times larger than that for two monomeric species (Table S6, calculated using the Smolu-chowski coagulation kernel^25^), and even more importantly, use of an oligomer-size dependent association rate does not affect calculation of nucleation time (Section S2.3). An oligomer size independent dissociation rate can be explained for a model where any binding interface consists of only two protein molecules. This is supported by observation of an increase in fibrillation rate on sonication that is expected to increase the number of fibril ends.^26^ Figure 3A shows that *C*_F_ has a typical S-shaped temporal profile wherein there is a long induction time during which *C*_F_ ≈ 0. This induction time, defined as aggregation nucleation time *t*_nu_, was calculated as the time at which *C*_F_ is equal to 5 % of its (long time) plateau value. The monomer and dimer concentrations obtained using the reduced kinetic model are in excellent agreement with the full model up to this nucleation time. The reduced kinetic model is clearly not appropriate for *t* ≥*t*_nu_, and therefore, *C*_F_ calculated using the reduced model does not have the S-shape profile, but still shows the presence of an induction time. In fact, the estimate of *t*_nu_ obtained from the two kinetic models are in excellent agreement, and this is not dependent on protein concentration and the method used to estimate the nucleation time (Section S2). This small effect of fibrillation can be explained on basis of very small fibril concentration till the nucleation time. Therefore, we conclude that the reduced kinetic model can be used to accurately determine the aggregation nucleation time, and the monomer and dimer concentration up to *t*_nu_. This forms a very important basis for our approach to estimate physically correct values for various rate constants in the kinetic model.

**Figure 1:**
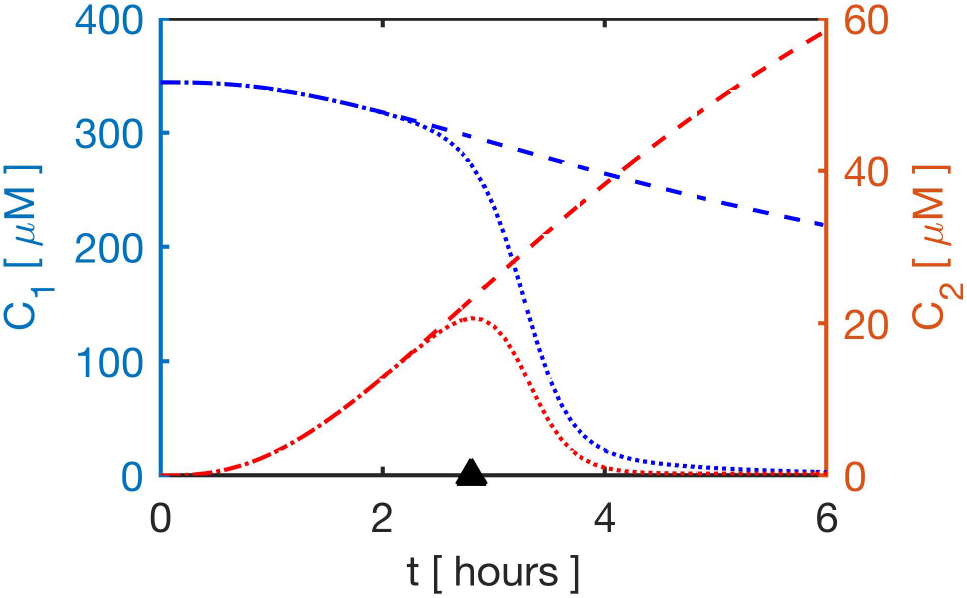
Reduced kinetic model for aggregation nucleation. Monomer (*C*_1_, in blue on left axis) and dimer (*C*_2_, in red on right axis) concentrations obtained by solving either the full kinetic model (*··*) or the reduced kinetic model (- -). Both models were solved using the kinetic parameters reported in Murray et al ^12^ at 344 µM insulin. The aggregation nucleation time (▴) is shown on time axis.

### Diffusional Association Rates

Under the assumptions made here, the reduced kinetic model is fully characterized by thermodynamics and kinetics of two events: (i) protein folding-unfolding and (ii) protein dimerization involving at least one aggregation prone species (*A*). We have taken insulin folding (*k*_f_) and unfolding (*k*_u_) rates from our previous study where metadynamics simulations were used to characterize the insulin folding landscape.^27^ Further, three metastable basins, each consisting of kinetically connected partially-folded conformations (collectively labeled as *A* here), in addition to a native (*N*) and a denatured (*D*) basin were identified in that study. A first-principles treatment of addition of *A* to a monomer (*N* or *A*) or a sub-nuclei oligomer (*A*_*i*_, 2 ≤*i* ≤ 5), as given in Table 1, requires the determination of rate constants (*k*_a_, *k*_d_) for reversible diffusional association of the *N*.*A* complex. For the association rate *k*_a_, we have used the transition complex theory that has been shown to provide accurate predictions for kinetics of diffusional association of several protein-receptor pairs in their native state.^28,29^ Specifically, it was shown to suitably account for the effect of long-range electrostatic interactions on the so-called basal association rate, the latter determined solely from unbiased translation and rotation of a pair of rigid-bodies to form native protein-protein contacts. This is important in the present case since there is a net charge of +7 *e* (at pH 2) on the *N*.*A* dimer. Given the structural heterogeneity of partially-folded species *A* and lack of knowledge on the structure of transient *N*.*A* dimer, we first determined an ensemble of representative structures for the *N*.*A* dimer to be used as input to the transition complex theory.

### Structure of N.A dimer(s) in optimal binding mode

A set of 16 most stable (lowest binding free energy) *N*.*A* insulin dimer conformations were taken from our earlier study wherein extensive rigid-body docking was used to generate a large ensemble of possible dimer conformations, followed by conformation ranking on basis of binding free energies calculated using the molecular mechanics Poisson-Boltzmann surface area (MM-PBSA) implicit solvent model.^30^ We carried out an extensive set of molecular dynamics simulations starting with these 16 structures with a specific aim of allowing re-orientation of binding partners under influence of long-range electrostatics and refinement of contacts determined from rigid-body docking. To allow for the determination of an electrostatically optimal binding mode, five replicate simulations were done for each rigid-body docked structure wherein monomer *A* in any given *N*.*A* dimer was moved perpendicular to the binding interface by a distance equal to twice the Debye length (2*λ*_D_ =2.2 nm at 100 mM NaCl) to ensure a negligible effect of any specific protein-protein interactions (PPIs) at this initial separation (see Figure S1). Since at this step we are only interested in capturing the effect of long-range electrostatic interactions between the *N* and *A* monomers screened by the aqueous buffer, we have used the coarsegrained (CG) MARTINI forcefield with the polarizable water model.^31^ This was shown to accurately reproduce screening effects in aqueous solutions, and therefore, provides best compromise between speed and accuracy: all-atomistic simulations spanning several microseconds for a system with over one million atoms are intractable while implicit solvent forcefields (for example, G ō -like models) will not correctly reproduce screened electrostatic interactions. Each CG simulation was run for 300 ns, giving a total of 24 µs simulation time.

41 out of 80 CG-MARTINI simulations resulted in re-formation of the *N*.*A* dimer for an extended time during the 300 ns simulation. In all cases where a complex was formed, it was found to have a stable conformation for at least last 100 ns of the simulation (Figure S3). However, the binding interface area for the final structure obtained after CG-MARTINI simulation was found to be lower than that for the rigid-body docked dimer (Table S2), possibly resulting from a larger bead size in the CG model. We therefore carried out a 20 ns all-atomistic (AA) simulation for each of the 41 re-formed complexes, with the initial structure taken as the central structure of the highest population cluster in the stable part of the CG simulation. The binding interface area quickly increased during the AA simulation to a value very close (in most cases greater) to that in the rigid-body docked dimers (Table S2 and S3) and, thereafter, remained stable (data not shown). For each of the 41 *N*.*A* dimers obtained at the end of AA simulations, the highest binding interface similarity score (equation S13) with any one of the 16 rigid-body docked complexes lies between 0.39–0.74 (maximum possible similarity score value is 1), thus indicating a refinement in dimer structure rather than a complete change. In summary, the combination of CG-MARTINI and AA simulation allowed us to obtain a set of optimized dimer structures, with the former allowing us to achieve a better electrostatic complementarity at a manageable computational cost and the latter providing for better packing at the binding interface.

### Kinetically Dominant Binding Mode(s)

From the large set of binding modes for formation of a *N*.*A* dimer, we expect only a small subset to be relevant for aggregation nucleation. To find this subset of important binding modes, we first group kinetically similar dimer structures together using a reaction criterion employed in Brownian dynamics simulation schemes for diffusional protein-protein association: a successful encounter complex was considered to be formed when two independent binding interface contacts were formed simultaneously.^32^ We obtained eight clusters on applying this criterion in-combination with integer linear programming to minimize the number of clusters (details in Section S1.2). Each cluster (largest: 16 structures, smallest: 2 structures), consisting of structures with a distinct set of binding interface residues is representative of a distinctive binding mode (full list of cluster sizes and binding modes in Figure S4).

We solved the subset of rate equations in **Scheme B** up to the formation of the *N*.*A* complex to determine which binding modes dominate the formation of this diffusional dimer. The calculation of diffusional association rate (*k*_a_) using the transition complex theory requires the existence of a transition state separating the bound and the unbound state of a complex where no large-scale conformational changes have occurred. Figure 2A shows that indeed the transition state of *N*.*A* dimer is located at the outer boundary of the bound state, the latter defined by a deep interaction energy well. As done in the original work, the total number of contacts in any configuration, *N*_c_, has been used as a measure of interaction energy. Figure 2B shows a sharp increase in standard deviation of the rotational angle (*χ*), *σ*_*χ*_, at the contact level of the transition state 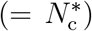, indicative of a large increase in rotational freedom on reaching the transition state. In fact, 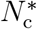 was determined from an order parameter based on the distribution *σ*_*χ*_(*N*_c_). For several associating protein pairs with unlike charges, it was shown that the transition state involved the formation of an electrostatically complementary conformation that lead to an enhancement of protein association rates compared with the basal association rate.^28,32^ Table 2 shows that indeed (*k*_a_) for various *N*.*A* insulin dimers is strongly correlated with the electrostatic interaction energy of the transition state complex, 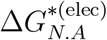, whereas only weakly correlated with the binding free energy of the bound complex, 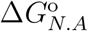. The Pearson correlation coefficient for top five binding modes (as per (*k*_a_)) was obtained to be −0.83 for the former and −0.57 for the latter. In spite of the same sign of net charge on *N* (+4*e*) and *A* (+3*e*) monomers, Δ*G*_elec_ has a small positive value (∼thermal energy (= 1.5*RT* = 1 kcal mol^−1^)) for dimers with four highest (*k*_a_). Overall, we have shown that the key requirements concerning the nature of the transition complex postulated in the theory by Zhou and co-workers, so-far used only for modeling native protein-protein association, are satisfied in the present case for non-native association leading to aggregation nucleation. The estimation of the dissociation rate, *k*_d_, from the binding free energy of *N*.*A* dimer, 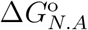, provided us the full set of kinetic parameters (given in Table 2) required to obtain the dimer concentration in each binding mode.

**Figure 2:**
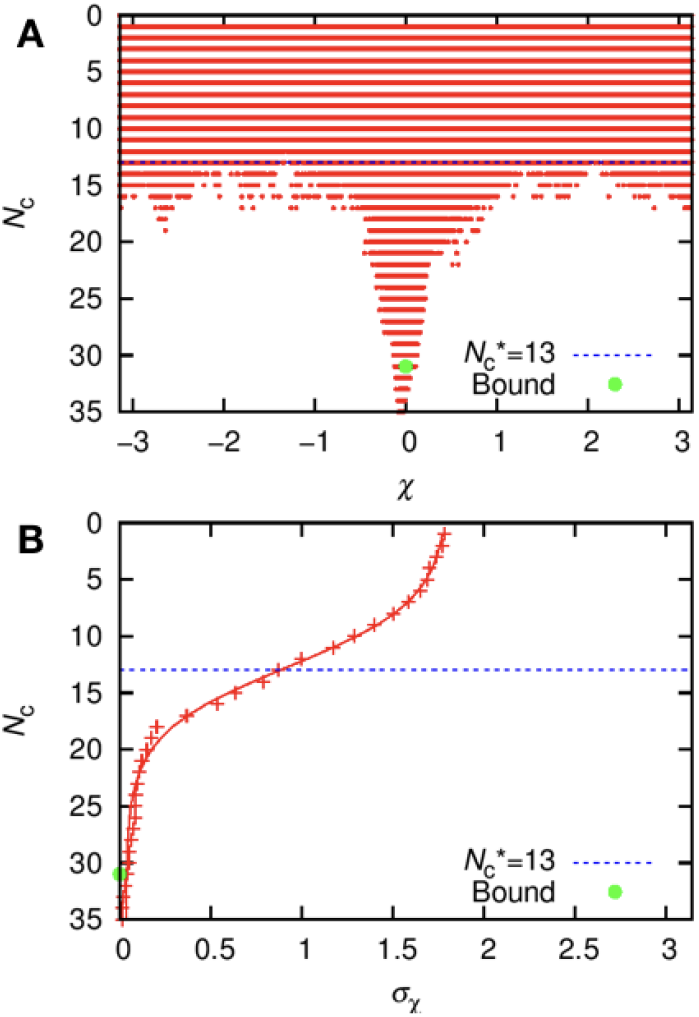
Energy landscape for *N*.*A* complex. **(A)** Scatter plot for total number of contacts, *N*_*C*_, in the ensemble of structures obtained on a rotation *χ* away from the bound complex. **(B)** Standard deviation of the rotational angle, *σ*_*χ*_, plotted versus *N*_*C*_. The data is shown for the kinetically dominant *N*.*A* dimer. The total number of contacts in the bound state and in the transition state was 32 and 13, respectively. Plots generated using TransComp. ^33^

**Table 2:** Kinetic and thermodynamic parameters for eight binding modes of *N*.*A* dimer. The association rate of *N*.*A* dimer, *k*_a_, and the electrostatic interaction energy of the corresponding transition state complex, 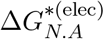, were determined using the transition complex theory. The binding free energy of the *N*.*A* dimer, 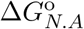, was calculated using the MM-PBSA method on a 20 ns MD trajectory. It also includes the contribution from protein configurational entropy change estimated using an atom specific weighted surface area (WSAS) method. ^34^

| $NA^i$ | $k_a^i$<br>($\mu M^{-1} hr^{-1}$) | $\Delta G_{N.A}^{*(elec)}$<br>kcal mol$^{-1}$ | $\Delta G_{N.A}^o$<br>kcal mol$^{-1}$ |
| --- | --- | --- | --- |
| 1 | 6734 | 0.694 | -3.84 |
| 2 | 1332 | 0.937 | -1.97 |
| 3 | 536 | 1.237 | -2.99 |
| 4 | 376 | 1.184 | 0.234 |
| 5 | 350 | 1.498 | -2.85 |
| 6 | 147.2 | 3.221 | 1.2 |
| 7 | 12.08 | 5.123 | 0.64 |
| 8 | 4.06 | 1.781 | -2.24 |

**Figure 3:**
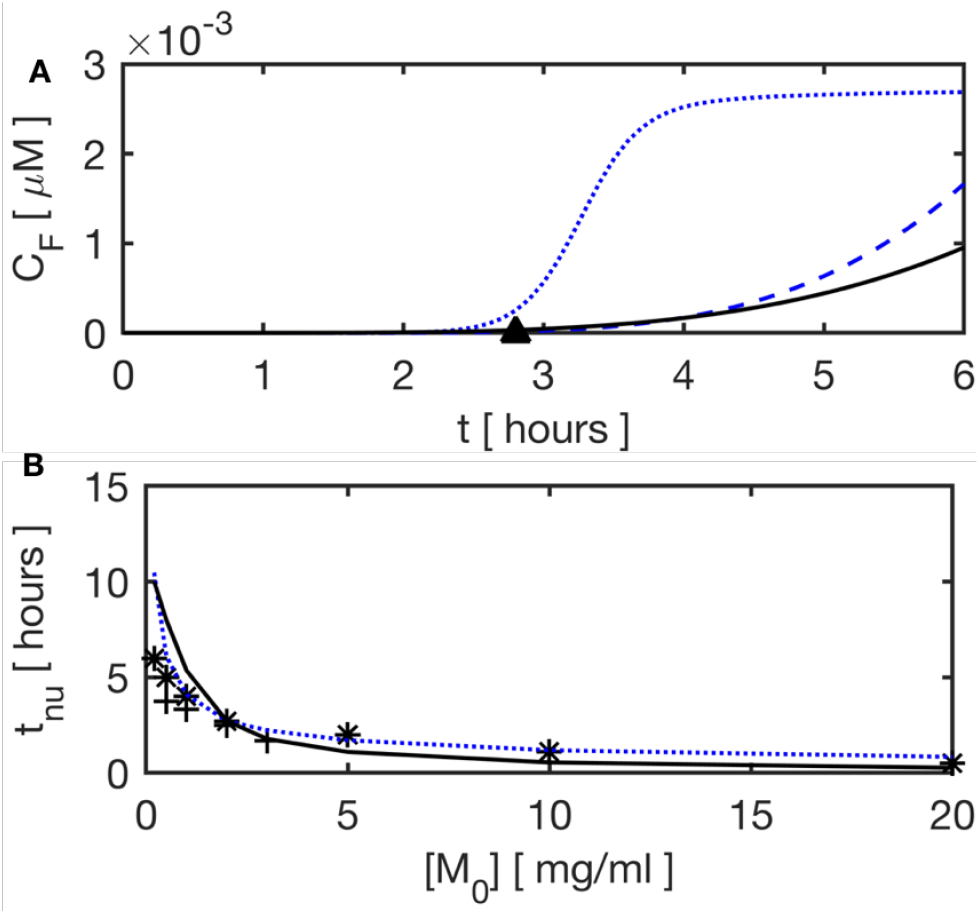
Effect of protein (insulin) concentration on aggregation kinetics. **(A)** Temporal evolution of fibril concentration, *C*_F_, obtained using the full kinetic model (*··*), and the reduced kinetic models either excluding (- -) or including (—) dimer conformational rearrangement. **(B)** Aggregation nucleation time (*t*_nu_) as function of initial insulin concentration ([*M*_0_]) obtained from kinetic models compared with the corresponding experimental estimate ((*\**) Lee et al. ^16^, (+) Amdursky et al. ^55^).

For each of the eight binding modes, the structure with the lowest (most negative) 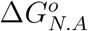 (preferable over 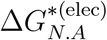 since it iscorrelated with both *k*_a_ and *k*_d_) was used for the determination of temporal evolution of the concentration of *N*.*A* dimer ([*N*.*A*]) in that mode. Table 2 shows there is significant variation in *k*_a_ and 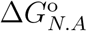 for these eight structures.

The *k*_a_ values of order 10^3^ µM h^−1^ obtained here are typical for diffusional association of protein complexes (e.g., see Table 3), but are seven orders of magnitude larger that the over-all association rate, *k*_nu_, for the formation of the final dimeric complex, *A*_2_ (Table 3). This clearly indicates that insulin dimerization is not diffusion-limited and must involve a slow conformational rearrangement step, as given by equations 5 and 6 in **Scheme B**. We discuss one important implication of this revised kinetic scheme in the later part of this letter where it is shown that inclusion of conformational rearrangement is necessary for quantification of effect of ligands on insulin aggregation nucleation time. Figure S5A shows that one particular binding mode (binding mode 1 in Table 2) has a significantly higher value for [*N*.*A*] compared to any other mode. We acknowledge that this observation of a single dominant binding mode analogous to native protein-receptor binding might be fortuitous resulting from approximations involved in calculation of binding energy using the MM-PBSA-WSAS approach. A direct validation of 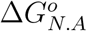 estimated here is not possible as corresponding experimental estimation of equilibrium data for non-native insulin dimerization is made difficult by presence of. However, the value obtained (−3.84 kcal mol^−1^ for the most stable binding mode in Table 2) is comparable to that for native insulin dimers, which is the dominant species at acidic pH.^17^ For example, the binding free energy for dimerization of porcine insulin (different from human insulin only at amino acid 30 of B chain: TB30A) at 298 K and pH 2, determined using concentration difference spectroscopy, was reported to be −6.8 kcal mol^−1^.^35^ The presence of specificity in non-native interactions can also be related to the observation that the partially-folded, aggregation-prone conformations of insulin and many other proteins are known to have a distinct structure with residual native contacts.^36,37^ Therefore, we expect to see similarities in characteristics of protein association involving such PFIs and that involving only native proteins, as observed also in applicability of the transition complex theory to the *N*.*A* dimer. We, therefore, conjecture that only a small number of binding modes, even if not only one mode as determined here, will be relevant for insulin aggregation nucleation and subsequent fibrillation. The presence of structural specificity in protein interactions involved in insulin aggregation is also supported indirectly by some experimental data, such as the proposal that fibril growth occurs by addition of an insulin PFI only to the ends of an existing fibril,^38^ significant inhibition of insulin aggregation at equimolar or smaller concentrations of a ligand with high affinity for insulin PFIs.^39,40^ The identification of dominant binding modes has allowed us to determine the hotspot residues and dominant interactions in non-native association of insulin (Figure S5B) that can in-turn be used as input for high-throughput screening of ligands for disrupting protein-protein interactions leading to insulin aggregation.

**Table 3:** Kinetic parameters in insulin aggregation nucleation. Column **A**: Present work; Column **B**: Full set of parameters determined empirically by fitting to aggregation growth data; ^12^ Column **C**: direct experimental measurements for individual elementary steps for insulin or other globular proteins.

| Rate constant | A | B | C |
| --- | --- | --- | --- |
| $k_{\text{nu}}, \mu\text{M}^{-1} \text{ h}^{-1}$ | $1.59 \times 10^{-3}$ | $1.8 \times 10^{-4}$ | $(10^{-3} - 10^{-4})^{41}$ |
| $K_{A_2}, \mu\text{M}$ | 22 | 45 | $12^{35,42}$ |
| $k_a$ | $6.74 \times 10^3$ | - | $(10^3 - 10^6)^{28,43,44}$ |
| $K_{N.A}, \mu\text{M}$ | $3.3 \times 10^3$ | - | - |
| $f$ | .07 | .99 | $(0 - 0.15)^{27}$ |
| $k_u, \text{h}^{-1}$ | $2.4 \times 10^8$ | 0.81 | $(10^5 - 10^{11})^{45,46}$ |
| $k_f, \text{h}^{-1}$ | $2.8 \times 10^9$ | $3.6 \times 10^{-3}$ | - |
| $k_r, \text{h}^{-1}$ | 5.2 | - | $0.1^{26,47}$ |

### Role of Dimer Conformational Rearrangement

We have noted above a significant difference in diffusional association rate for *N*.*A* dimer (*k*_a_ *∼* 10^3^ µM h^−1^) and the estimate for association rate of final dimeric species *A*_2_ (*k*_nu_ *∼* 10^−4^ µM h^−1^). This implies that *N*.*A* and *A*_2_ must be separated by a slow dimer conformational rearrangement step as shown in **Scheme B**(equations 4–6), with *k*_r_ representing the forward rate for conformational change of *N*.*A* to *A*_2_. Since *k*_nu_ *<< k*_a_, equation S26 simplifies to 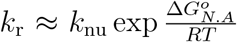, implying that a given overall dimerization rate, (*k*_nu_), can be obtained for infinite combinations of *k*_r_ and 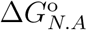. We have taken *k*_r_ as an adjustable parameter to be determined from data on aggregation growth while 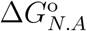 was calculated using the MM-PBSA-WSAS method as explained. In fact, *k*_r_ is the only adjustable parameter in our reduced kinetic model. All other kinetic parameters, reported in Table 3, were fixed at values obtained by our first-principles approach. The binding free energy of the aggregation pathway dimeric species (*A*_2_) is ex pected to be different from 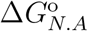 (because of dimer conformational rearrangement) and was taken to be equal to the experimental estimate available for binding free energy of native insulin dimerization (dimer equilibrium dissociasation constant *K*_*A*2_ = 12 µM corresponding to 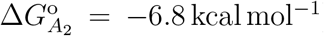. *k*_r_ was then estimated by solving the resulting reduced kinetic model **Scheme B** for time variation of total monomer concentration, *C*_1_(*t*), and performing a least-squares fit to *C*_1_(*t*) obtained from the full kinetic model without conformational rearrangement (**Schemes A, D**). The parameters for the full kinetic model were taken from an earlier study that showed excellent agreement with experimental data on fibril growth kinetics.^12^ We have used the data only up to the nucleation time (*t*_nu_ = 2.8 h at 344 µM insulin) in-accordance with the validity of reduced kinetic model and obtained *k*_r_ = 5.2 h^−1^. A simulated annealing protocol with several distinct initial guesses for *k*_r_ gave the same solution in each case. Even when *K*_*A*2_ was also taken as an unknown parameter, we obtained *k*_r_ = 5.05 h^−1^, essentially the same as previous estimate. In this case, we obtained 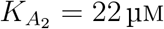, very close to the fixed value of 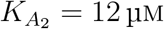. This implies that *k*_r_ = 5.2 h^−1^ is a robust estimate for rearrangement rate and is several orders of magnitude smaller than effective diffusional association rate (*k*_a_*C*_1_(0) = 2.3 *×* 10^6^ h^−1^). Slow rearrangement of pre-fibrillar aggregates has been reported for insulin^48^ and other small globular proteins,^47,49,50^ with rates several orders of magnitude smaller than those corresponding to protein folding or *β*-sheet formation at interface of two small peptides. The proposal that this much smaller rearrangement rate compared to intra- and inter-protein *β*-sheet formation can be explained on basis of double layer repulsion and reconfiguration^51^ is not supported by the observations in our study. For example, for an initial separation of more than two Debye lengths between two associating insulin monomers (*N* and *A*) ensuring almost complete formation of a ionic double layer with a total protein charge of +7 *e*, 50 % of stable dimer complexes (total 80 simulations) were formed in less than 100 ns of simulation time. An alternate mechanism, based on characterization of early events of insulin aggregation using orthogonal bioanalytical techniques, proposes a slow, sequential rearrangement of pre-fibrillar oligomeric species after diffusional association.^48,52^ This slow rearrangement of prefibrillar aggregates on a timescale of more than 10 h has also been reported for other amyloid proteins such as *Aβ*40.^47,49,50^

Using the estimate for *k*_r_ in equation S26, we get the overall dimer association rate *k*_nu_ = 1.6 × 10^−3^µM^−1^ h^−1^, which is an order of magnitude larger than the value obtained when all kinetic rate constants were treated as adjustable parameters and determined simultaneously using least-squares fitting^12^ (see Table 3). This difference in *k*_nu_ is offset by the difference in the native monomer fraction for folding-unfolding equilibria: we have used a value of *f* = 0.07 as obtained from metadynamics simulations^27^ in contrast to *f* = 0.99 obtained by least-squares fitting.^12^ Insulin formulations incubated at aggregation-prone conditions of acidic pH (2–3) and high temperature (40 °C– 60 °C), but low concentration (≈ 100 nm), had the native protein as dominant species as determined using Fourier transform infrared (FTIR) spectroscopy with *f* = 0–0.15.^53,54^ Therefore, simultaneous estimation of multiple kinetic parameters by fitting to experimental data on growth kinetics can yield estimates significantly different from values obtained by direct quantitative methods.

### Prediction of Effect of Protein and Ligand Concentration on Aggregation Nucleation

Figure 3A shows the comparison of prediction for stable oligomer concentration (*C*_*i*≥6_ *≡C*_F_) obtained by solving the full kinetic model (parameters from ref.^12^), and the reduced kinetic model without rearrangement (parameters from ref.^12^) or with rearrangement (present work, parameters in Table 3). All three cases show almost equal induction time. However, as discussed above, the reduced model does not have an inflection point in *C*_F_(*t*), and therefore, can not be used to directly determine the aggregation nucleation time, *t*_nu_. Instead, we used the relative fibril concentration at *t*_nu_, defined for a given initial protein concentration [*M*_0_] as 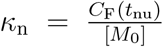, calculated at 344 µM insulin as a fixed parameter to determine the nucleation time at all other state points(Section S2.2 for details). Even when *κ*_n_ was estimated independently at each [*M*_0_], it stays essentially constant for large changes in [*M*_0_] ((2-20) mg ml^−1^) (Table S5). The monomer and dimer concentrations obtained using either of the two reduced models are in excellent agreement with the full model up to *t*_nu_ (Figure S9), implying that both reduced models can be used to obtain estimates for aggregation nucleation time.

We first compare predictions from different kinetic models and parameter estimation methods against the data available on effect of initial insulin concentration on aggregation nucleation time. We have kept all model parameters, estimated either by fitting to aggregation growth data at 344 µM (1.97 mg ml^−1^) insulin (*k*_r_, *κ*_n_) or by direct determination for the elementary step under consideration (*k*_u_, *k*_f_, *k*_a_, *K*_*N*.*A*_, *K*_*A*2_), fixed at all protein and ligand concentrations (given in Table 3). Figure 3B shows that estimate for *t*_nu_ obtained using these physically determined parameter values in the reduced kinetic model are in excellent agreement with corresponding values determined from a diverse set of bioanalytical techniques for initial insulin concentrations in range (0.1 to 20) mg ml^−1 16,55^ leading to a variation of over two orders in *t*_nu_. The full kinetic model, using parameters determined by least-squares fit to aggregation growth data at 344 µM,^12^ provides predictions for *t*_nu_ at the same accuracy level. In both cases, a higher estimate (compared to experimental data) for *t*_nu_ at low concentrations could be explained by increased importance of heterogeneous nucleation (from contained walls, solution-air interface) at low concentrations that has not been included in our kinetic scheme.

We observe that large differences in kinetic parameter values obtained using direct determination at the level of elementary step versus the case when full set is determined empirically (see Table 3) does not affect the quality of presdiction of aggregation kinetics variation with protein concentration. This can be attributed to absence of any competing reaction steps in the kinetic schemes when investigating role of protein concentration, thereby making possible offsetting of deviations in model parameter values. For example, a significantly larger value for the equilibrium fraction of aggregation-prone monomer was offset by a reduced dimerization rate in the empirically determined set. A more stringent test is provided by cases where conditions are changed to affect a particular elementary step (for instance, a sequence mutation that affects protein stability), affect various elementary steps differently (for instance, change in salt concentration will have different effect on folding equilibria and association equilibria), or when a competing reaction is added to the kinetic model. Here we have taken up the last case, wherein a ligand with specific affinity for the aggregation-prone species (*A*) was added to the protein formulation.

We have used the data available on inhibition of insulin aggregation by 1,2-Bis[4-(3-sulfonatopropoxyl)phenyl]-1,2-diphenylethene salt (BSPOTPE), an organic fluorogen that was postulated to bind predominantly at hydrophobic sites on the aggregation-prone monomeric and oligomeric species.^39^ In it the photo luminescence (PL) spectrum was used to determine aggregation nucleation time for insulin incubated with BSPOTPE at 330 K and pH 2. The additional kinetic equations to account for ligand (*L*) effect are given by **Scheme C** (equation 7). For calculation of kinetic parameters, several putative structures for the *A*.*L* complex were determined by separately docking five distinct conformations of BSPOTPE at each of the ten dimerization hotspot residues of partially folded species *A*. The top docked structures were found to be metastable with the binding interface staying essentially the same over a 20 ns MD simulation (for example see Figure S6A and S6B). The binding interface area of species *A* in the most stable *N*.*A* complex 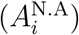 and in the most stable *A*.*L* complex 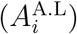, shown in Figure 4(inset), was 4.1 nm^2^ and 2.8 nm^2^, respectively. This implies that only one BSPOTPE molecule can bind to the hotspot residues of *A* in the *N*.*A* complex, and accordingly, our reaction scheme involved only one ligand binding event. The *A*.*L* structure with highest cos Θ^*A*.*L,N*.*A*^ (defined in equation S13), a parameter calculated from *A*.*L* binding energy and similarity of binding interface residues in this complex to insulin dimerization hotspot residues, was used for the calculation of ligand association rate from the transition complex theory (see Methods Section S1.7 for details). We obtained *k*_aL_ = 9.23×10^4^ µM^*−*1^ h^*−*1^, which is an order of magnitude higher than the diffusional association rate of *N*.*A* complex and can be explained on basis of smaller size of the ligand compared to native insulin and a smaller binding interface area for the *A*.*L* complex compared to the *N*.*A* complex. The equilibrium dissociation constant for the *A*.*L* complex (*L*: BSPOTPE), *K*_A.L_, was taken to be 10 µM, as calculated from the photo luminescence (PL) spectrum for BSPOTPE solution with fibrillar insulin.^39^ In the same study, no emission was detected for the solution with native insulin, and therefore, our reaction scheme does not involve ligand binding to native insulin. Overall, **Scheme C** for ligand inhibition was solved with the schemes with no insulin dimer intermediate with the empirical parameter set (Table 1 **Scheme A+D** or **Scheme A** only) or the scheme with a dimeric intermediate with parameter set determined in this work (Table 1 **Scheme B**). The insulin concentration was kept fixed at 500 µM and BSPOTPE concentration was varied between (0 to 500) µM in accordance with the experimental study. Figures 4, S7(B) shows that the schemes with no dimeric intermediate completely fail to predict the effect of a ligand with specific affinity for the partially folded species. The inclusion of the intermediate provides very accurate estimate for *t*_nu_ over the entire ligand concentration range with nucleation time varying over two orders of magnitude, viz. from a few hours in absence of ligand to more than a week (175 h) at equimolar ligand concentration. This data clearly indicates the important role of competition between formation of an on-pathway dimeric intermediate involving species *A* leading to aggregation and inhibition of this pathway by a small molecule ligand with high affinity for the hot spot residues of *A* involved in dimer formation. The *A*.*L* complex has a stability comparable to the insulin dimer, viz. *K*_A.L_ = 10 µM versus 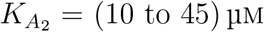, but a significantly higher association rate. Even a one order of magnitude change in *K*_A.L_, at fixed *k*_aL_, gives essentially the same *t*_nu_ (Figure S7(A)), indicating that within the range studied ligand association rate has a more important role than the ligand dissociation rate in prediction of aggregation nucleation time. Further, this implies that free energy estimation methods that provide reasonably accurate estimates for protein-ligand binding free energy^56^ can be combined with the approach presented here to provide a fully predictive tool for accurate estimation of ligand effects on protein aggregation inhibition.

**Figure 4:**
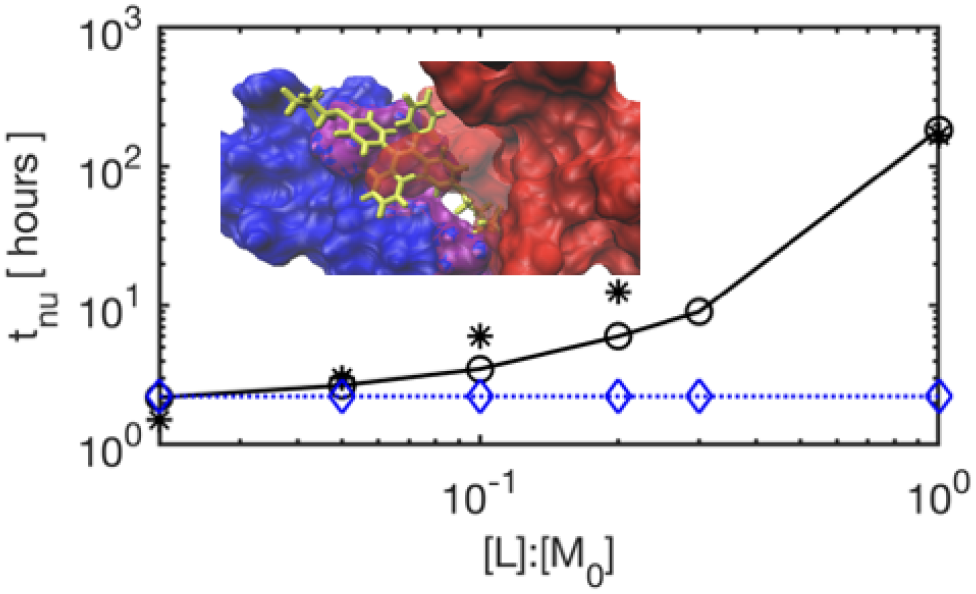
Effect of ligand (BSPOTPE) concentration on aggregation kinetics. Aggregation nucleation time (*t*_nu_) as function of BSPOTPE concentration (*L*) at 500 µM insulin ([*M*_0_]) obtained by solving either the full model with empirical parameter set from ref. ^12^ (–♦–) or the reduced model with parameters determined in this work (– ○ –). In both cases, ligand inhibition was modeled using equation 7. Experimental estimate from Hong et al. ^39^ is also shown (∗). Inset: A snapshot of the overlay of kinetically dominant *N*.*A* complex and the *A*.*L* com-plex zoomed-in at the binding interface (*N*: red (surf view), *A*: blue (surf view), *L*: green (bonds view).

To summarize, we have determined a set of transferable parameters for extended Lumry-Eyring aggregation kinetics of insulin using a combination of techniques for modeling elementary events spanning several orders of timescales (milliseconds to hours) while still accounting for specific inter-molecular interactions: insulin folding kinetics from all-atomistic metadynamics simulations (prior work), an aggregation pathway dimeric intermediate from structural bioinformatics and multiscale molecular dynamics simulations, and dimer association rates from the transition com-plex theory. Interestingly, we found that association kinetics for kinetically dominant modes for formation of an aggregation-prone insulin dimer have characteristics very similar to that of native protein association such as a transition state at the outer boundary of a deep and narrow bound state ensemble. A single set of physically meaningful kinetic parameters was shown to provide very accurate estimates for changes in aggregation nucleation time from less than a hour to over a week with changes in either insulin or the ligand (BSPOTPE) concentration. We expect these conclusions to be valid in general for other globular proteins. Finally, we note that the approach presented here can be directly applied to quantitatively estimate effect of small molecule ligands on aggregation inhibition of globular proteins, thereby, greatly aiding in development of high stability therapeutic formulations using high-throughput computational screening methods. Development of computational and experimental approaches for determination of kinetics and thermodynamics of non-native association involving slow conformational rearrangement can further the application of kinetic modeling to protein aggregation.

## Methods

### Diffusional Association Complex

The all atomistic (AA) *N*.*A* complex was moved perpendicular to the binding interface by a surface separation of 2*λ*_*b*_ (*λ*_*b*_: Debye length) and was then converted into MARTINI coarsegrained (CG) representation. The short-ranged non-bonded interactions between CG beads were scaled down since earlier studies have shown that the MARTINI forcefield over- estimates protein-protein interactions.^57^ The reduced interaction strength was chosen such that more than half of rigid-body docked *N*.*A* dimers, found to be metastable in earlier all-atomistic simulations,^30^ stay in an associated state with a stable binding interface during CG simulation (see details in Table S1). CG-MD simulations were carried out using GROMACS at aggregation-prone experimental conditions of 338 K, pH 2, and 600 µM insulin. Elastic restraints on backbone of both metastable species (*N*&*A*) were implemented using the ELNEDYNE algorithm.^58^ The charge state of the titrable residues (HIS, LYS, ARG) and the end terminals was fixed according to their *pK*_a_ (obtained using the *propka* server^59^). Details of the protocol to obtain a total charge of +7 *e* on the dimer, in accordance with ion mobility mass spectroscopy measurements, are provided in our previous work.^30^ The final structure from CG-MD simulations was refined by backmapping to the AA representation followed by a 20 ns MD run using the CHARMM36 force-field.^60^ The central structure of the largest cluster from the stable portion of this AA MD trajectory was taken as the diffusional association complex. The details of simulation parameters, equilibration protocol, and validation of simulation convergence for AA and CG MD simulations are provided in Section S1.A.1 and S1.A.2. The diffusional association rate for a unique set of *N*.*A* dimers (see Section **Linear Integer Programming**) was calculated using the transition complex theory^28^ (See S1.3). To this end, we used the *Transcomp* server^33^ that takes as input the structure of the bound complex and the ionic strength (set to 100 mM in accordance with simulations and other calculations in this study).

### Linear Integer Programming (LIP)

The obtained *N*.*A* dimers were classified into a smallest number of clusters such that structures in the same cluster have similar polar contacts while in different clusters have sufficiently distinct contacts. A set of unique polar contacts at the binding interface was defined using the criterion applied for formation of an encounter complex in Brownian dynamics simulations^61^ (see Figure S4B). The optimization problem to determine the smallest number of clusters with a central structure, referred to as the parent structure, assigned to each cluster was solved using LIP with following set of constraints: (1) any two parent structures should have less than two common polar contacts, (2) any structure with ≥2 common polar contacts with all structures in a cluster was assigned to that cluster. Full details on implementation of LIP are given in Section S1.2

### Binding Energy

Standard-state binding energy for the *N*.*A* dimer and the *A*.*L* complex, Δ*G*°, was calculated using the implicit-solvent molecular mechanics Poisson-Boltzmann surface area (MM-PBSA) method. We have used the single trajectory approach, shown to have smaller standard error than the multiple trajectory approach, wherein averages for both the apo (unbound) and holo (in complex) states are calculated from the MD trajectory of only the solvated complex.^62^ A trajectory of 20 ns was used for both the complexes. The individual contributions to Δ*G*°, as given by equation S8, were obtained using the *g mmpbsa* utility.^63^ We have also added the configurational entropy contribution to Δ*G*°, which was calculated using a set of atom-specific, empirically determined weights for the buried and the solvent-accessible area of the atoms at the binding interface of given complex.^34^ The Δ*G*° thus obtained is referred to as MM-PBSA-WSAS binding energy. Specific details on individual terms and the parameter values used in calculation are given in Section S1.4.

### Population Balance

The set of ordinary differential equations (ODEs) for various kinetic schemes (given in Section S3) was solved using MATLAB ®. The range of parameter values used here resulted in a set of stiff ODEs and accordingly, we used the ode23tb solver (uses a combination of trapezoidal rule and backward difference formula) with a small time step of 0.2 h. Other stiff equation solvers, such as ode15s and ode23s, lead to large deviations from requirement of total protein mass conservation. For the case of unknown kinetic parameter determination, we used the lsqnonfit function in-combination with ode23tb. Within this the total squared error between the target and the predicted monomer species concentration up to the nucleation time *t*_nu_ was minimized. MultiStart function was used wherein the minimization problem is solved by starting from multiple start points and its output consists of all local solutions identified ranked from best (lowest objective function value) to worst (highest objective function value).

### Ligand Association

Several conformations for the *A*.*L* complex were generated using *Hex 8*.*0*.*0* ^64^ *by rigid-body docking of each of the five stable conformations of BSPOTPE (L*) reported in Hong et al.^39^ onto centroids defined by each of the ten residues of *A* identified as binding hotspot residues in the dominant *N*.*A* complex. 1000 unique structures were generated for each set (for given centroid and given ligand conformation) and the most stable *A*.*L* complex, as per the scoring function in *Hex*, for each ligand conformation was selected. The 10 structures thus obtained were solvated at 100 mM NaCl and a 20 ns MD trajectory was generated for each complex. The ligand topology was obtained using the *cgenff* server.^65^ To get the representative *A*.*L* complex for the calculation of association rate, first the central structure of the largest cluster was selected from the stable portion of each MD trajectory. Then the structure with the highest cos Θ^*A*.*L*;*N*.*A*^ (defined in Equation S13) was used as input to the *Transcomp* server for calculation of *k*_aL_. Additional details in section S1.7.

## Acknowledgement

This work was funded and supported by the Science & Engineering Research Board, Department of Science & Technology, GoI (EMR/2017/004218) and Ministry of Human Resource Development, GoI fellowship. The authors thank IIT Delhi HPC facility for computational resources.

## Supporting Information Available

Additional methodology details: AA and CG simulation setup, docking and binding energy calculations, linear integer programming, population balance method and comparison of *t*_nu_ calculated through different methods.

