## supplementary information. for "A Multiscale Model for Quantitative Prediction of Insulin Aggregation Nucleation Kinetics"

### Multiscale Predictive Modeling of Insulin Aggregation Nucleation Kinetics

### Supporting Information Available

### S1 Methods

#### S1.1 Diffusional Association

##### S1.1.1 Coarse Grained (CG) Molecular Dynamics (MD)

*Plane fitting.* The interface residues in the native protein $N$ were identified and a best fit plane was found using an in-house MATLAB code as shown in Supplementary Information (SI) Figure S1. The normal to the plane is the direction vector along which the $A$ is shifted by a distance of twice Debye length ($2\lambda_d = 2.2\,\text{nm}$)

*ELNEDYNE Restraint.* The separated dimer, was converted into MARTINI CG structures with elastic restraints on backbone of $N$ and $A$ in the metastable $N.A$ complex using the ELNEDYNE algorithm.[1] Most CG forcefields are biased towards native contacts which makes them unsuitable for study of partially folded protein diffusion. ELNEDYNE algorithm enables us to put elastic restraints on the secondary structures. In SI Fig S2 , we track the RMSD of the backbone $C_\alpha$ atoms of the partially folded proteins and native insulin identified

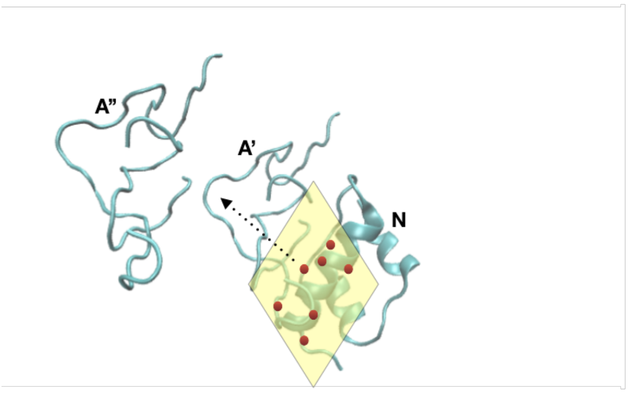


**Figure S1: Plane Fitting on interface:** Plane fitted to the $C_\alpha$ atoms of the interface residues of the native protein,$N$. The aggregation prone protein $A$ moved along the normal by a distance of Debye length showing its new position $A''$

in our previous work.[2] RMSD $< 0.35\,\mathrm{nm}$ suggests minimal change in secondary structure over the length of simulation of 300 ns.

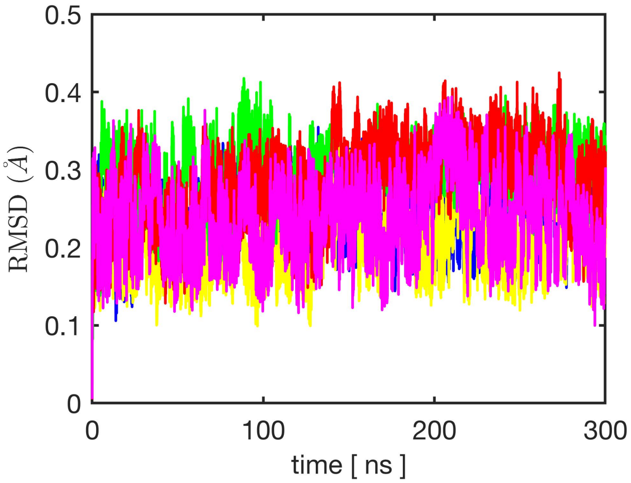


**Figure S2: Stability of secondary structures of $PF_i$:** RMSD of the $C_\alpha$ backbone atoms of the native(yellow) , $\mathrm{PF}_1$(pink), $\mathrm{PF}_2$(red) and $\mathrm{PF}_3$(green) after running a simulation with MARTINI force field with ELNEDYNE elastic restraints on the secondary structures of the complex.

*Re-scaling MARTINI.* The short-ranged non-bonded interactions between CG beads were scaled down as earlier studies have shown that the MARTINI forcefield overestimates the protein-protein interactions.[3] The reduced interaction strength was chosen such that more than half of rigid-body docked $N.A$ dimers, found to be metastable in earlier all-atomistic simulations,[4] stay in the associated state for a large portion of the CG simulation. The scaling was done using a scaling parameter, $\alpha$, with $\alpha$ =1 and $\alpha = 0$ corresponding to the use of the original MARTINI force field and the weakest possible MARTINI interaction respectively, for all Van der Waals interactions. The value $\alpha = 0$ is the so-called 'repulsive' interaction, for which MARTINI sets the well-depth to 2.0 kJ/ mol. For a given value of $\alpha$, the value of the well-depth, $\epsilon$ in the Lennard Jones 12-6 potential function, applied to any particular interaction, is determined using

$$\epsilon = \epsilon_{\text{repulsive}} + \alpha \times (\epsilon_{\text{original}} - \epsilon_{\text{repulsive}}) \tag{S1}$$

where, $\epsilon_{\text{repulsive}}$ is 2.0 kJ/mol, and $\epsilon_{\text{original}}$ is the original MARTINI well-depth used for the interaction of interest. Different values of $\alpha$ was used to study the stability of the insulin $N.A$ complex. We scaled down the parameter until 50 % of the total simulations form a stable complex. The percentage of the structures that formed a stable complex for various scaling factors has been listed in SI Table S1.

**Table S1:** As MARTINI over estimates the short range interactions, the forcefield parameters were scaled down. The scaling down parameter $\alpha$ was determined by running a MD simulation for 100 ns on each of the 16 top ranked complexes obtained from Avinash et al.[4] and calculating $S_{\%}$,the percentage of total simulations in which the starting complex remained stable for at least 50 ns. The scaled down well-depth, $\epsilon$, in Lennard Jones 12-6 potential function for a scaling factor of $\alpha$ is given by $\epsilon = \epsilon_{\text{repulsive}} + \alpha \times (\epsilon_{\text{original}} - \epsilon_{\text{repulsive}})$ where, $\epsilon_{\text{repulsive}}$ is 2.0 kJ/mol, and $\epsilon_{\text{original}}$ is the original MARTINI well-depth used for the interaction of interest

| $\alpha$ | $S_{\%}$ |
|---|---|
| 0.65 | 18.5 % |
| 0.70 | 26.25 % |
| 0.75 | 37.5 % |
| 0.80 | 51.25 % |

We find that when the short range interactions values were scaled down to 80 % of their actual value, 51 % structures remain associated. The short range interaction parameters in the MARTINI force field were scaled down to 80 % of the actual values and each top ranked structure was simulated independently five times resulting in total eighty simulations.

*Simulation Protocol and Parameters.* CG-MD were carried out using GROMACS 5.1.4

at aggregation-prone experimental conditions of 330 K , pH 2, and 600 µM of insulin. The pH of the solution was fixed by altering the charges on the titrable residues (HIS, LYS, ARG) and end terminal of $A$ depending on the $pK_a$ obtained using the *propka* server[5]). Details of the method to fix the protonation state has been outlined in our previous work.[4]

The converted protein complexes were first energy minimized in vacuum with full protein restraint and then solvated in a cubic box of 18 nm and a salt concentration of 100 mM (typical for insulin formulations) and energy minimized again to remove bad contacts. Solvation is done using MARTINI polarizable water model. The whole system contained $\approx$ 45000 CG polarizable water molecules and the protein dimer which had 1575 atoms is reduced to 232 atoms in the CG model. The minimization were done using steepest descent algorithm.

The solution was then equillibrated using NVT ensemble in two steps to bring it to the desired temperature of 330 K . First with a full position restraint on all the atoms of the protein for 100 ps followed by restraint only on the backbone atoms, using a v-rescale thermostat with a low temperature coupling constant, $\tau_T$ = 0.5 ps. The system was then equillibrated in a NPT ensemble using Parrinello-Rahman barostat for 500 ps, with pressure coupling constant of $\tau_P$ = 12 ps and with a new $\tau_T$ = 6 ps. Both v-rescale thermostat and Parrinello-Rahman barostat are recommended to be used for MARTINI (CG) force field. The final equillibrated system was then run for 300 ns with the NPT ensemble as a production run. The long range interaction were modelled using reaction-field for all the runs. Clustering was done on the stable part of the trajectory on the basis of root mean square deviation (RMSD) of the $C_\alpha$ backbone atoms with a cut-off of 0.35 nm. The central structure of the most populated cluster obtained was considered the final structure. We track the $\cos\Theta^{N.A;N.A}$ (refer equation S13 but no scaling by $\overrightarrow{G^L}$) i.e inner product between $A$ in final $N.A$ complex obtained from simulation and $A$ in the complexes formed during simulation. We report the values for all parent structures. All the complexes show convergence during the course of simulation and the same is observed for all trajectories in which a complex is obtained (data not shown) .

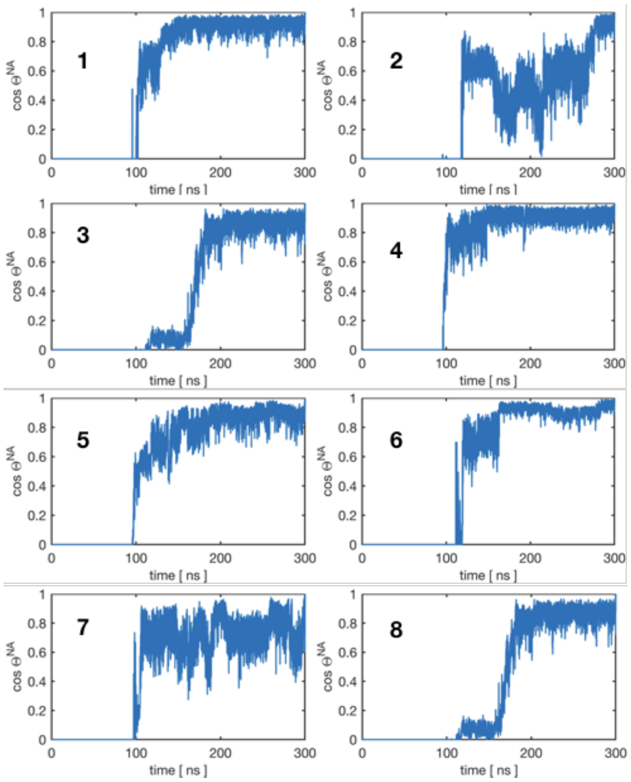


**Figure S3: Convergence of MD trajectories:** cos $\Theta^{N.A;N.A}$ tracked for every frame during the CG-MD trajectory for all the parent structures is reported showing a stable value for at least 200 ns

##### S1.1.2 All Atomistic (AA) MD

The final structure from Section S1.1.1 was back mapped to its all AA structure and the system is set for another MD run using the CHARMM36 forcefield.[6] The equillibration protocols followed for CG simulation in Section S1.1.1 were applied for AA simulation also. The final structure from CG simulation were solvated in a box by placing the protein complex in centre of a cubic box and fixing a minimum distance of 1.5 nm from the edge resulting a box size of 8 nm . The system had ≈ 17000 molecules including water , protein and salt ions. Equillibration is done with NVT ensemble with full protein restraint for 100 ps followed by a run with restraint only on the backbone with NVT ensemble for 100 ps followed by NPT ensemble with no restraint for 500 ps and finally moving onto the production run for 20 ns to allow for the secondary structure fluctuation and better packing of interface. A $\tau_T =$ 0.1 ps and $\tau_P = 2$ ps was used during equillibration. For the production run $\tau_T = 15$ ps and

0.5 ps for protein and non protein was used respectively, and $\tau_P = 4$ ps was set. The central structure of the largest cluster after analysis of the AA MD trajectory is the final diffusional association complex.

The structures obtained from CGMD were extended to AAMD to enhance the packing and allow for secondary structure fluctuation. It is evident from Table S2 and Table S3 which shows improvement in the interface area and inner product respectively and further reinforces our choice of extending our CG simulations by a small AA simulation.

**Table S2:** Interface area of the structures obtained at the end of CG run and after extending it in its back-mapped AA form (in $\mathrm{nm}^2$). The structures in bold are the parent structures.

| Nos | CG | AA | Nos | CG | AA | Nos | CG | AA |
|---|---|---|---|---|---|---|---|---|
| **1** | **2.2** | **3.1** | 15 | 2.8 | 3.1 | 29 | 4.0 | 5.9 |
| **2** | **6.2** | **7.7** | 16 | 4.2 | 5.8 | 30 | 5.0 | 8.4 |
| 3 | 5.3 | 7.1 | **17** | **4.4** | **5.7** | 31 | 5.0 | 6.7 |
| 4 | 4.5 | 6.9 | 18 | 4.8 | 6.5 | 32 | 6.3 | 7.5 |
| 5 | 6.1 | 8.2 | 19 | 4.4 | 6.5 | 33 | 5.3 | 7.6 |
| 6 | 3.8 | 5.4 | 20 | 6.9 | 9.1 | 34 | 6.4 | 7.7 |
| 7 | 4.2 | 5.6 | 21 | 5.0 | 8.7 | **35** | **4.5** | **7.7** |
| **8** | **5.0** | **7.1** | 22 | 5.9 | 7.8 | 36 | 4.7 | 5.9 |
| 9 | 3.2 | 4.1 | **23** | **4.0** | **5.9** | 37 | 2.6 | 3.9 |
| 10 | 5.0 | 7.7 | 24 | 3.2 | 5.1 | 38 | 4.8 | 7.9 |
| 11 | 4.8 | 6.8 | 25 | 5.3 | 7.1 | 39 | 6.3 | 8.3 |
| 12 | 4.8 | 6.4 | **26** | **3.5** | **4.1** | 40 | 1.6 | 3.7 |
| 13 | 4.8 | 6.9 | 27 | 4.1 | 4.5 | 41 | 6.6 | 8.3 |
| 14 | 3.8 | 5.6 | **28** | **3.4** | **6.9** | | | |

We also report the highest inner product, $\cos\Theta^{N.A;N.A}$ for the 16 top ranked docked structure against 1) set of diffusional complex obtained from its own set of simulation (ip1) 2) the set of all diffusional complexes (ip2) (Table S4). We observed that ip2 has a significantly high value (at least $\geq 0.5$) which shows our diffusional complexes have shown good enough sampling to show good similarity with metastable docked structures.

**Table S3:** Inner product, $\cos\Theta^{N.A;N.A}$ of the structures obtained at the end of CG run and after extending it in its back-mapped AA form against the initial docked structure. The structures in bold are the parent structures.

| Nos | CG | AA | Nos | CG | AA | Nos | CG | AA |
|---|---|---|---|---|---|---|---|---|
| **1** | **0.44** | **0.63** | 15 | 0.51 | 0.73 | 29 | 0.46 | 0.66 |
| **2** | **0.24** | **0.35** | 16 | 0.39 | 0.57 | 30 | 0.26 | 0.38 |
| 3 | 0.69 | 0.61 | **17** | **0.12** | **0.17** | 31 | 0.43 | 0.62 |
| 4 | 0.15 | 0.22 | 18 | 0.67 | 0.45 | 32 | 0.15 | 0.25 |
| 5 | 0.45 | 0.65 | 19 | 0.18 | 0.26 | 33 | 0.63 | 0.61 |
| 6 | 0.42 | 0.60 | 20 | 0.64 | 0.62 | 34 | 0.56 | 0.60 |
| 7 | 0.27 | 0.38 | 21 | 0.15 | 0.22 | **35** | **0.52** | **0.74** |
| **8** | **0.19** | **0.14** | 22 | 0.26 | 0.37 | 36 | 0.56 | 0.77 |
| 9 | 0.12 | 0.21 | **23** | **0.60** | **0.47** | 37 | 0.26 | 0.38 |
| 10 | 0.29 | 0.42 | 24 | 0.44 | 0.64 | 38 | 0.43 | 0.61 |
| 11 | 0.12 | 0.18 | 25 | 0.12 | 0.18 | 39 | 0.40 | 0.57 |
| 12 | 0.50 | 0.72 | **26** | **0.30** | **0.42** | 40 | 0.37 | 0.53 |
| 13 | 0.25 | 0.37 | 27 | 0.50 | 0.52 | 41 | 0.19 | 0.27 |
| 14 | 0.58 | 0.74 | **28** | **0.24** | **0.34** | | | |

**Table S4:** The binding interface area of top 16 docked structures is listed. ip1 is the highest $\cos\Theta^{N.A;N.A}$ of docked structure against set of diffusional complex obtained from its own set of simulation. ip2 is the highest $\cos\Theta^{N.A;N.A}$ of docked structure against the whole set of all diffusional complexes.

| Nos | Area ($nm^2$) | ip1 | ip2 | Nos | Area ($nm^2$) | ip1 | ip2 |
|---|---|---|---|---|---|---|---|
| 1 | 6.32 | 0.63 | 0.63 | 9 | 5.55 | 0.47 | 0.47 |
| 2 | 4.34 | 0.65 | 0.65 | 10 | 6.63 | 0.64 | 0.64 |
| 3 | 4.69 | 0.38 | 0.45 | 11 | 5.12 | 0.42 | 0.69 |
| 4 | 6.82 | - | 0.51 | 12 | 6.91 | 0.52 | 0.56 |
| 5 | 4.73 | 0.72 | 0.72 | 13 | 5.74 | 0.66 | 0.66 |
| 6 | 5.15 | 0.74 | 0.74 | 14 | 4.11 | - | 0.27 |
| 7 | 6.81 | 0.45 | 0.47 | 15 | 6.81 | 0.38 | 0.77 |
| 8 | 6.45 | 0.62 | 0.62 | 16 | 6.37 | 0.57 | 0.61 |

#### S1.2 Linear Programming

The obtained $N.A$ dimers were classified into a smallest number of clusters such that structures in the same cluster have similar polar contacts while in different clusters have sufficiently distinct contacts. A set of unique polar contacts at the binding interface was defined using the criterion applied for formation of an encounter complex in Brownian dynamics simulations[7] (see Figure S4). The optimization problem to determine the smallest number of clusters with a central structure, referred to as the parent structure, assigned to each cluster was solved using LIP with following set of constraints: (1) any two parent structures should have less than two common polar contacts, (2) any structure with $\geq 2$ common polar contacts with all structures in a cluster was assigned to that cluster. We found the minimum number of parent complexes that collectively cover all the possible binding modes available. It was achieved by converting the model into an optimization problem and using Linear programming to solve the model. The details of the equations for our optimisation models have been given below.

A matrix $A_{ij}$ is built by filling in with 1 and 0s. If the $i^{th}$ complex has more than 2 common independent contacts with the $j^{th}$ complex, then $A_{ij}$ is allotted 1 otherwise 0.

$$A = \begin{bmatrix} 1 & 0 & 0 & 0 & 1 \\ 0 & 0 & 1 & 1 & 0 \\ 1 & 1 & 0 & 1 & 1 \\ 0 & 0 & 0 & 0 & 1 \\ 1 & 0 & 0 & 0 & 0 \end{bmatrix} ; X = \begin{bmatrix} x_1 & x_2 & x_3 & ... \end{bmatrix} \tag{S2}$$

$X_i$ is an ensemble of structures which shows the interaction $i^{th}$ with other structures. We build the objective function to be minimized as AX = B

such that

$$B_i = \sum_{i=1}^{m} A_i X \tag{S3}$$

The following constraints are applied on the optimisation

$$B_i > 0 \tag{S4}$$

as each structure needs to have more than 2 common contacts with at least one complex(case where i=j). We also apply

$$.X_i \in \{0, 1\} \tag{S5}$$

where $X_i = 1$ means the row is considered in the solution else 0. `intlinprog` function of MATLAB is used to solve the optimization problem.

The resulting ensemble and its population has been listed in Figure S4. In case a structure had $\geq 2$ distinct common polar contacts with more than one ensemble, it is allotted the ensemble with which it shares more number of polar contacts.

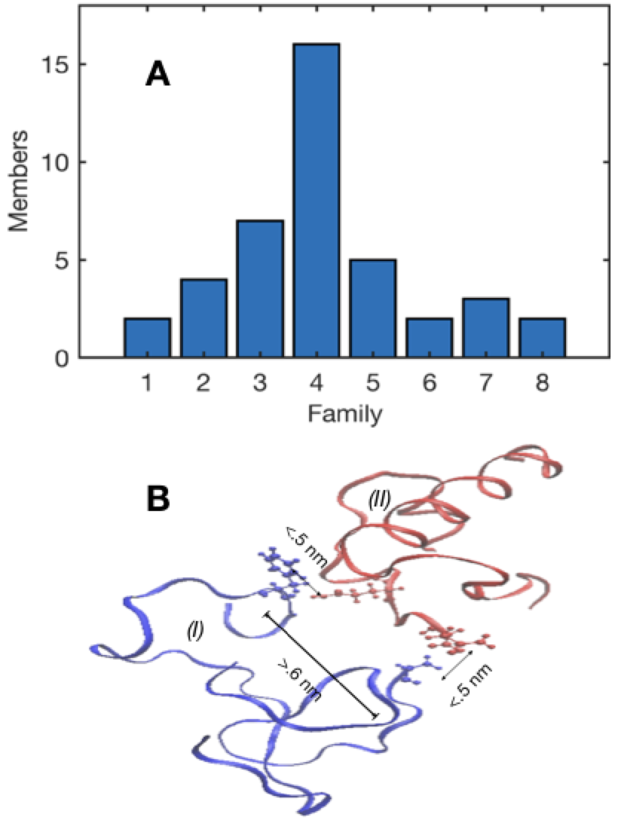


**Figure S4: Family of structures:** The complexes obtained at the end of MD simulations are categorized into families based on their common distinct polar "donor-acceptor" contacts on their interface according to the work of Gabdoulline et al.[8] Complexes belonging to the same family have two or more common polar contacts and one or zero common polar contacts with the complexes of a different family. **(A)** Distribution of obtained structures among various families. **(B)** The heavy atom inter-atomic distance between inter-protein residues needs to be less than $0.5\,\mathrm{nm}$ to be considered in contact and the distance between two intra-protein contact needs to be more than $0.6\,\mathrm{nm}$ to be considered as a distinct contact.

#### S1.3 Transition Complex Theory

The diffusional association rate between $A$ and $N$ was calculated using the transition complex theory.[9] The theory is based on the concept of a transition state, which lies between the bound and unbound state. The unbound state are guided by long range interactions to reach a transition state, and the numerous short- range interactions are enough to guide the transition state to the final complex. There is an instantaneous decrease in the translational and rotational freedom as the complex moves from bound state to transition state. This instantaneous change is manifested by the standard deviation of rotational angle of the protein,$\chi$ sampled at different contact levels (native + non-native),$N_c$. Equation S6 tracks the transition complex and is identified by the level of $Nc$ that maximizes

$$\Xi(N_c) =< \sigma_\chi(N_c') >_{N_c'<N_c} -\sigma_\chi(N_c) \tag{S6}$$

which represents the difference between the standard deviation of $\chi$ at contact level $N_c$ and the average for all lower contact levels $N_c'$. The transition complex obtained is used to calculate the association rate $k_a$ as

$$k_a = k_0 exp(-\Delta G_{\mathrm{elec}}/k_B T) \tag{S7}$$

where $k_0$ is the basal rate constant in absence of electrostatic interactions and $\Delta G_{elec}$ is the electrostatic enhancement. We use the *Transcomp* server[10] which takes in the bound complex as an input and allows us to set the ionic strength by giving the salt concentration as input which is 100 mM in our study

#### S1.4 Binding Energy

Standard-state binding energy for the $N.A$ dimer and the $A.L$ complex, $\Delta G^{\mathrm{o}}$, was calculated using the implicit-solvent molecular mechanics Poisson-Boltzmann surface area (MM-PBSA)

method. We have used the single trajectory approach, shown to have smaller standard error than the multiple trajectory approach, wherein averages for both the apo (unbound) and holo (in complex) states are calculated from the MD trajectory of only the solvated complex.[11] A trajectory of 20 ns was used for both the complexes. The individual contributions to $\Delta G^{o}$, as given by equation S8, were obtained using the *g_mmpbsa* utility.[12] We have also added the configurational entropy contribution to $\Delta G^{o}$, which was calculated using a set of atom-specific, empirically determined weights for the buried and the solvent-accessible area of the atoms at the binding interface of given complex.[13] The $\Delta G^{o}$ thus obtained is referred to as MM-PBSA-WSAS binding energy.

$$\Delta G^{o} = \Delta E_{\mathrm{MM}} + \Delta G_{\mathrm{solv}} - T\Delta S \tag{S8}$$

where

$$\begin{aligned}\Delta E_{\mathrm{MM}} &= \Delta E_{\mathrm{bonded}} + \Delta E_{\mathrm{non-bonded}}\\ &= \Delta E_{\mathrm{bonded}} + (\Delta E_{\mathrm{elec}} + \Delta E_{\mathrm{vdw}})\end{aligned}$$

and

$$\Delta G_{\mathrm{solv}} = \Delta G_{\mathrm{polar}} + \Delta G_{\mathrm{apolar}}$$

where $\Delta E_{\mathrm{bonded}}, \Delta E_{\mathrm{vdw}}, \Delta E_{\mathrm{elec}}$ correspond to the bond, angle, torsion, Van der Waals, and electrostatic terms in the molecular mechanical force field. The polar term, $\Delta \mathrm{G}_{\mathrm{polar}}$, was estimated by solving the Poisson–Boltzmann (PB) equation wherein the protein is modeled at all-atomic resolution and the solvent is taken as a continuum of known dielectric. The non-polar term, $\Delta \mathrm{G}_{\mathrm{apolar}}$, was estimated using the surface area model ($\Delta G_{apolar} = \gamma \Delta A$), where $\gamma$ is the surface tension associated with a hydrophobic interface. GROMACS 5.1.4 compatible *g_mmpbsa* package is used to calculate the MMPBSA values. Grid spacing of 0.51 Å is set. We have used a high solute dielectric of 4 as recommended for high charge species.[14] Solvent dielectric is fixed at 80. The probe radius for non polar energetic contribution was fixed 1.4 Å but a smaller probe radius of 0.6 Å was used as it shows better agreement with experimental values during dimer formation.[15]

The entropy was calculated as described in the methods reported by Wang et al[13] given in Equation S9 and S10

$$S = \sum_{i=1}^{N} w_i (SAS_i + k \times BSAS_i) \tag{S9}$$

$$BSAS_i = 4\pi (r_i + r_{\text{prob}})^2 \tag{S10}$$

Where, $w_i$ is the weight for atom i; $SAS_i$ is the solvent accessible surface area of atom i; N is the number of atoms in a molecule; $BSAS_i$, the buried solvent accessible surface area of atom i,is calculated using Equation S10 (note that BSAS of atom i is complementary to SAS of atom i); and $r_i$ is the radius of atom i. The probe radius, $r_{\text{prob}}$, was set to 0.08 nm. The term $k$ is the adjustable parameter with the values taken from the same work. In their approach, the conformational entropy of a molecule, $S$, can be obtained by summing up the contributions of all atoms, no matter they are buried or exposed. Each atom has two types of surface areas, solvent accessible surface area (SAS) and buried SAS (BSAS). Each area is weighted depending on the atom type involved with $k$ used as an adjustable parameter.

#### S1.5 Dominant Complex

To identify the kinetically dominant binding mode(s), we used the subset of rate equations up to the formation of this encounter complex, as given by equations S11 and S12:

$$\text{N} \underset{k_\text{f}}{\overset{k_\text{u}}{\rightleftharpoons}} \text{A} \tag{S11}$$

$$\text{N} + \text{A} \underset{k_\text{d}}{\overset{k_\text{a}}{\rightleftharpoons}} \text{N} \cdot \text{A} \tag{S12}$$

The diffusional association rate ($k_\text{a}$) for each $N.A$ dimer was determined using the transition complex theory,[9] For each of the eight binding modes, the structure with the lowest (most negative) $\Delta G^o_{N.A}$ (preferable over $\Delta G^{*(\text{elec})}_{N.A}$ since it is correlated with both $k_\text{a}$ and $k_\text{d}$ ) was used for the determination of temporal evolution of the concentration of $N.A$ dimer, $[N.A]$,

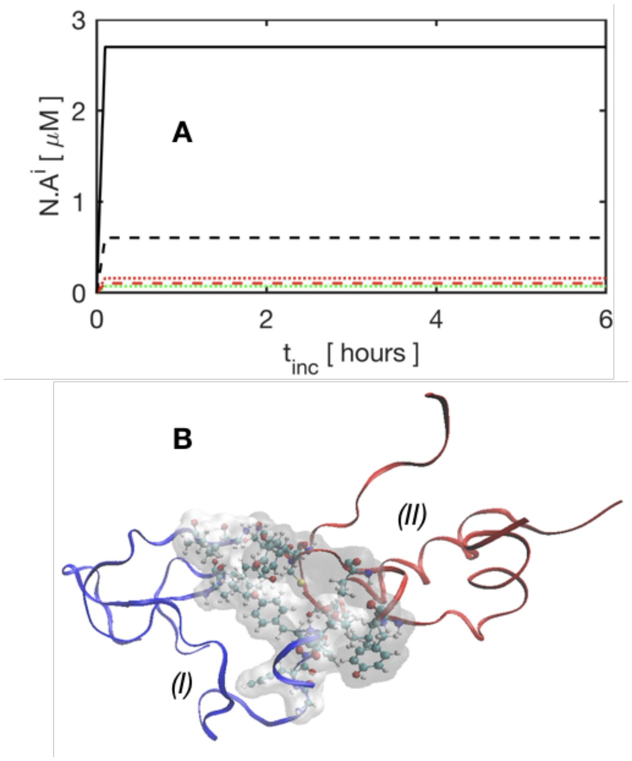


**Figure S5: Identification of dominant complex**. **(A)** The concentration profile of $[NA^i]$ obtained by solving the equations S11 and S12 using the parameters listed in Table 2 ,showing the existence of a dominant complex and its interface showing the hotspot residues for $A$ in blue *(I)* and $N$ in red *(II)* in panel **(B)**

in that binding mode. Figure S5A clearly shows the existence of a dominant complex. The identification of dominant binding modes has allowed us to determine the hotspot residues and dominant interactions in non-native association of insulin (Figure S5B). Total of 10 residues were involved in the partially-folded $A$ insulin namely $\mathrm{GLY}^{A1}$, $\mathrm{ILE}^{A2}$, $\mathrm{LEU}^{A13}$, $\mathrm{TYR}^{A14}$, $\mathrm{GLY}^{B44}$, $\mathrm{PHE}^{B45}$, $\mathrm{PHE}^{B46}$, $\mathrm{TYR}^{B47}$, $\mathrm{THR}^{B48}$ and $\mathrm{ALA}^{B51}$. Except $\mathrm{GLY}^{A1}$ and $\mathrm{ALA}^{B51}$, all the residues are uncharged and the interaction is mostly hydrophobic.

#### S1.6 Ligand Association

Several conformations for the $A.L$ complex were generated using *Hex 8.0.0*[16] by rigid-body docking of each of the five stable conformations of BSPOTPE ($L$) reported in Hong et al.[17] onto centroids defined by each of the ten residues of $A$ identified as binding hotspot residues in the dominant $N.A$ complex. 1000 unique structures were generated for each set (for given centroid and given ligand conformation) and the most stable $A.L$ complex, as per the scoring function in *Hex*, for each ligand conformation was selected. The 10 structures thus obtained were solvated at 100 mM NaCl and a 20 ns MD trajectory was generated for each complex. The ligand topology was obtained using the *cgenff* server.[18] To get the representative $A.L$ complex for the calculation of association rate, first the central structure of the largest cluster was selected from the stable portion of each MD trajectory. Then the structure with the highest $\cos\Theta^{A.L;N.A}$ (defined in Equation S13) was used as input to the *Transcomp* server for calculation of $k_{\mathrm{aL}}$. Each docked complex was placed inside a solvated box and same step wise equillibration protocol was followed as in Section S1.1.2 with the structure first being bought to a temperature of 330 K using NVT ensemble with full protein restraint and then moving onto NPT with partial restraints and finally NPT with no restraint on the docked structure with the same barostat, thermostat and coupling constants used as in section S1.1.2. The $A.L$ docked complex were simulated for 20 ns , during which the interface remain stable. The interface area, and the energy landscape of the top ranked $A.L$ complex has been shown in SI Figure S6D

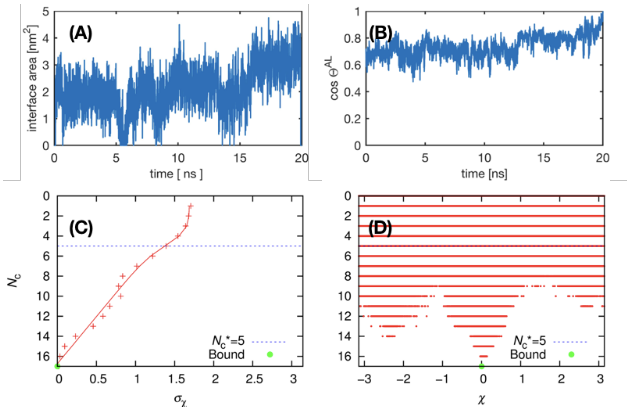


**Figure S6: Calculating $k_{aL}$: (A)** Interface area of the $AL$ complex during the course of simulation . **(B)** $\cos\Theta^{AL;AL}$ calculates the inner product between the $A$ of the final $AL$ complex and $A$ of the complexes formed during the trajectory **(C)** The standard deviation of the rotational angle $(\chi)$, $\sigma_\chi$, is plotted as a function of total number of contacts, $N_c$ for the $A.L$ complex. The transition state was determined from the distribution $\sigma_\chi(N_c)$. **(D)** Scatter plot for the total number of contacts, $N_c$, for the $A.L$ on a given rotation (specified by the rotational angle $\chi$) away from the bound complex. The data is shown for the highest ranked $A.L$ complex where the total number of contacts in the bound state and in the transition state was equal to 17 and 5, respectively.

Structures obtained from MD simulations were ranked on the basis of a weighted inner-product calculated from binding interface area of species $A$ in the $A.L$ complex and the selected $N.A$ complex, $\cos\Theta^{A.L;N.A}$, as defined in SI Equation S13.

$$\begin{aligned} \cos\Theta^{A.L;N.A} &= (\overrightarrow{G^L} \odot \overrightarrow{A^L}) \cdot \overrightarrow{A^P} \\ \text{where } G_i^L &= \frac{e^{\frac{-Ei}{kT}}}{\sum_{i=1}^{n} e^{\frac{-Ei}{kT}}} \end{aligned} \tag{S13}$$

$E_i$ is the contribution of $i^{th}$ residue of $A$ to the binding affinity of the $A.L$ complex and $n$ is the total number of residues of $A$, $\overrightarrow{A^L}$ contains residue-wise binding interface area for $A$ in the given $A.L$ complex, $\overrightarrow{A^P}$ contains residue-wise binding interface area for $A$ in the dominant $N.A$ complex. The $i^{th}$ element of $\overrightarrow{A^L}$ and $\overrightarrow{A^P}$ was calculated as

$$\begin{aligned} \overrightarrow{A_i^L} &= SASA_i^{\text{monomer}} - SASA_i^{\text{AL}} \\ \overrightarrow{A_i^P} &= SASA_i^{\text{monomer}} - SASA_i^{\text{NA}} \end{aligned}$$

where, $SASA_i^{\text{monomer}}$ is the solvent accessible surface area of the $i^{\text{th}}$ residue of $A$ in the unbound state, $\text{SASA}_i^{NA}$ is the solvent accessible surface area of the $i^{th}$ residue of $A$ in the $N.A$ complex, $\text{SASA}_i^{AL}$ is the solvent accessible surface area of the $i^{th}$ residue of $A$ in the $A.L$ complex . The binding interface area for $i^{\text{th}}$ residue in $A.L$ complex was multiplied by the Boltzmann weight obtained from its binding energy contribution, as represented by the Hadamard product ($\odot$) between $\overrightarrow{A^L}$ and $\overrightarrow{G^L}$. Then, the inner product between the resultant vector and $\overrightarrow{A^P}$ was used as the ranking score.

We also observe no major change in $t_{\text{nu}}$ when $k_{dL}$ was varied by a order of magnitude showing less sensitivity towards the parameter $k_{dL}$ making our model very robust(refer Fig S7A).

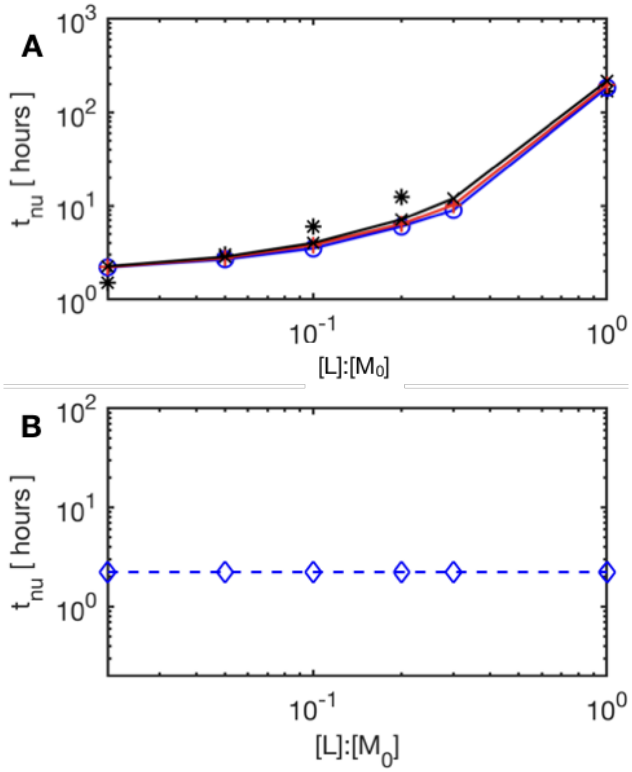


**Figure S7: Effect of ligand on insulin aggregation nucleation time. (A)** Nucleation time $t_{\text{nu}}$ calculated for three different $K_{AL}$ values: ( $- \circ -$ ) $K_{AL} = 10\,\mu\text{M}$ , $(- + -)$ $K_{AL} = 5\,\mu\text{M}$ and $(- \times -)K_{AL} = 1\,\mu\text{M}$. ( $*$ ) represents data from Hong et al.[17] **(B)** Nucleation time calculated using the reduced kinetic model with parameter values for protein folding and various protein-protein association steps taken from Murray et al.[19] and parameters for protein-ligand association-dissociation kinetics were taken from the present study.

### S2 Nucleation time

#### S2.1 Point of Inflection Method

The fibril concentration,$C_F$, profile of a typical protein aggregation process,**Scheme A**, is a sigmoidal curve as shown in Figure S8. The curve contains the initial exponential phase, then a small approximate linear phase and a final asymptotic phase. The linear phase of the curve represents the nucleation phase. In the present work, the time at which $C_F$ reaches 5% of its asymptote value is considered $t_{\text{nu}}$ and find $t_{\text{nu}} = 2.8\,\text{hours}$ for an initial $[M_0] = 344\,\mu\text{M}$.

$t_{\text{nu}}$ can also be defined as the time when the curve transits from exponential phase to the asymptotic phase. We find the inflection point of the curve and draw a tangent at the point. The point where the tangent meets the time axis, is considered as the $t_{\mathrm{nu}}$, and find $t_{\mathrm{nu}}$= 2.8 hours. We find no significant difference in $t_{\mathrm{nu}}$ from both the methods.

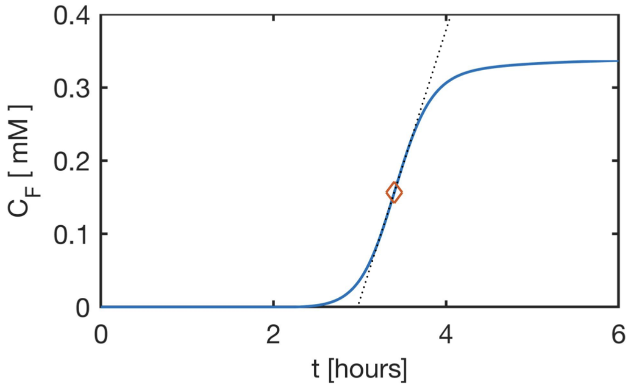


**Figure S8: Obtaining $t_{nu}$ from sigmoidal curve of fibril concentration,$C_F$:** Concentration profile of $C_F$ for a $[M_0] = 344\,\mu\mathrm{M}$ with the tangent at the point of inflection shown.

#### S2.2 Relative Fibril Concentration Method

As we do not model the fibrillation part of aggregation, we do not get a sigmoidal curve. We calculate the relative concentration $\kappa_{\mathrm{n}} = \frac{C_{\mathrm{F}}(t_{\mathrm{nu}})}{[M_0]}$ where $[M_0]$ is the initial monomer concentration for which $t_{\mathrm{nu}}$ is being calculated. We used $[M_0] = 344\,\mu\mathrm{M}$ to calculate $\kappa_{\mathrm{n}}$ but we show below in Table S5 the choice of $[M_0]$ to calculate $\kappa_{\mathrm{n}}$ has negligible effect on the final $t_{\mathrm{nu}}$. We track $\kappa_{\mathrm{t}} = \frac{C_{\mathrm{F}}(t)}{[M_0]}$ during aggregation and the time where $\kappa_{\mathrm{n}} = \kappa_{\mathrm{t}}$ is reported as $t_{\mathrm{nu}}$. We list the values of $\kappa_{\mathrm{n}}$ calculated at various $[M_0]$ and find no significant change in their value. The range of $t_{\mathrm{nu}}$ calculated for $344\,\mu\mathrm{M}$ lied between 2.4 - 3.3 for various $\kappa_{\mathrm{n}}$ reported in Table S5

**Table S5:** $\kappa_n$ at various initial monomer concentration, $[M_0]$

| $[M_0][mg/ml]$ | $\kappa_n(\times 10^{-8})$ | $t_{nu} for 344\,\mu M [hours]$ |
|---|---|---|
| .2 | 9.82 | 3.3 |
| .5 | 4.25 | 3.1 |
| 1 | 8.31 | 3.2 |
| 2 | 2.32 | 2.7 |
| 3 | 2.12 | 2.4 |
| 5 | 2.98 | 2.9 |
| 10 | 3.30 | 3 |
| 20 | 3.32 | 3 |

#### S2.3 Smoluchowski Kernel Method

A system whose entities count is tracked using PB method, the rates can be predicted using a Smoluchowski Kernel as shown in Equation S14

$$k_{i,j} = \frac{k_b}{W} B_{i,j} P_{i,j} \tag{S14}$$

where $k_b$ is the Smoluchowski rate constant for diffusion limited aggregation, defined as follows: $k_b = 8\mathrm{kT}/3\eta$, where k is the Boltzmann constant, T is the temperature, and $\eta$ is the viscosity. W is the so-called stability ratio, accounting for the energetic barrier between two approaching primary particles, $B_{i,j}$ is the correction for the Brownian term that accounts for the fractal nature of the aggregates and for the aggregation between unequal size clusters.

$$B_{i,j} = \frac{(i^{\frac{1}{D_f}} + j^{\frac{1}{D_f}})(\frac{1}{i^{\frac{1}{D_f}}} + \frac{1}{i^{\frac{1}{D_f}}})}{4} \qquad P = (ij)^{\lambda} \tag{S15}$$

where the experimentally measured fractal dimension of $D_f = 2.6$ has been used and semi-empirical parameter $\lambda = 0.61$[20] We have assumed a single kinetic rate upto the nucleation step. We re-solved the Scheme 2 using the rates obtained the Smoluchowski kernel(refer Table S6) and recalculated the nucleation time using the method described in Section S2.2. The $t_{nu}$ was found to be 2.3 hours.

**Table S6: Smoluchowski kernel rates:** Higher order oligomer association rates $k_{nu}^{ij}$(rate of association of $A_i$ and $A_j$) in terms of $k_{nu}^{11}$ as $k_{nu}^{ij} = \alpha \times k_{nu}^{11}$

| $k_{nu}^{ij}$ | $\alpha$ |
|---|---|
| $k_{nu}^{12}$ | 1.46 |
| $k_{nu}^{13}$ | 1.87 |
| $k_{nu}^{14}$ | 2.15 |
| $k_{nu}^{15}$ | 2.25 |

### S3 Modified Kinetic Rates

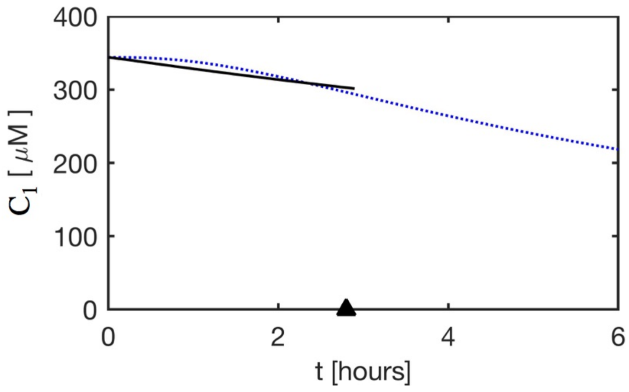


**Figure S9: Calculation of $k_r$ by fitting monomer data**: $k_r$ in Equation S26 was used as a fitting parameter and fitted to the monomer data of Murray et al. upto the nucleation time to obtain the values. Goodness of fit has been shown in the figure with ( ·· ) showing Murray et al monomer data without fibrillation and ( – ) showing monomer data in present study upto $t_{\text{nu}}$ .

$$\mathrm{N} + \mathrm{A} \underset{k_{-nu}}{\overset{k_{nu}}{\rightleftharpoons}} \mathrm{N_2}$$

$$\mathrm{N} + \mathrm{A} \underset{k_d}{\overset{k_a}{\rightleftharpoons}} \mathrm{NA} \underset{k_{-r}}{\overset{k_r}{\rightleftharpoons}} \mathrm{N_2}$$

$\mathrm{M} = \mathrm{N} + \mathrm{A}$ where M is the total Monomer concentration and $\mathrm{A} = \mathrm{f[M]}$ where f is the fraction of the total monomer concentration as partially folded A.

Kinetic differential equations for reaction scheme A.

$$\frac{dN}{dt} = -k_{nu}[N][A] + k_{-nu}[N_2] \tag{S16}$$

$$\frac{dA}{dt} = -k_{nu}[N][A] + k_{-nu}[N_2] \tag{S17}$$

$$\frac{dN_2}{dt} = k_{nu} f(1-f)[M]^2 - k_{-nu}[N_2] \tag{S18}$$

Adding equation (S16) and (S17)

$$\begin{aligned} \frac{d(N+A)}{dt} &= -2k_{nu}[N][A] + 2k_{-nu}[N_2] \\ \implies \frac{d(M)}{dt} &= -2k_{nu} f(1-f)[M]^2 + 2k_{-nu}[N_2] \end{aligned} \tag{S19}$$

Kinetic differential equations for reaction scheme B.

$$\frac{dN_2}{dt} = k_r[NA] - k_{-r}[N_2] \tag{S20}$$

$$\frac{dN}{dt} = -k_a[N][A] + k_d[NA] \tag{S21}$$

$$\frac{dA}{dt} = -k_a[N][A] + k_d[NA] \tag{S22}$$

adding equation (S21) and (S22)

$$\begin{aligned} \frac{dM}{dt} &= -2k_a[N][A] + 2k_d[NA] \\ \implies 2k_d[NA] &= \frac{dM}{dt} + 2k_a f(1-f)[M]^2 \end{aligned} \tag{S23}$$

substituting $\frac{dM}{dt}$ from equation (S19) into equation (S23)

$$\begin{aligned} 2k_d[NA] =& -2k_{nu} f(1-f)M^2 + 2k_{-nu}[N_2] \\ &+ 2k_a f(1-f)M^2 \\ \implies [NA] =& \frac{1}{k_d}[f(1-f)M^2(k_a - k_{nu}) \\ &+ k_{-nu}[N_2]] \end{aligned} \tag{S24}$$

substituting equation (S24) in equation (S20)

$$\frac{dN_2}{dt} = [f(1-f)\frac{k_r}{k_d}(k_a - k_{nu})][M^2] + \left[\frac{k_r k_{-nu}}{k_a} - k_{-r}\right][N_2] \tag{S25}$$

Comparing equations (S25) and (S18) we have

$$k_{nu} = \frac{k_a k_r}{k_r + k_d} \quad ; \quad k_{-nu} = \frac{k_d k_{-r}}{k_r + k_d} \tag{S26}$$
